# Generalized mathematical model of cancer heterogeneity

**DOI:** 10.1101/2020.05.25.115725

**Authors:** Motohiko Naito

## Abstract

The number of reports on mathematical modeling related to oncology is increasing with advances in oncology. Even though the field of oncology has developed significantly over the years, oncology-related experiments remain limited in their ability to examine cancer. To overcome this limitation, in this study, a stochastic process was incorporated into conventional cancer growth properties to obtain a generalized mathematical model of cancer growth. Further, an expression for the violation of symmetry by cancer clones that leads to cancer heterogeneity was derived by solving a stochastic differential equation. Monte Carlo simulations of the solution to the derived equation validate the theories formulated in this study. These findings are expected to provide a deeper understanding of the mechanisms of cancer growth, with Monte Carlo simulation having the potential of being a useful tool for oncologists.

## Introduction

Experimental oncology has significantly progressed over the last two decades (1,2). In particular, the discovery of cancer stem cells and the accumulation of knowledge on intratumor heterogeneity represent milestones in these progressions (3,4). Cancer stem cells appear to have common properties among different cancers (1), such as leukemia (3,5), brain cancer (6,7), and colorectal cancer (8,9), as does intratumor heterogeneity (2).

The number of studies related to mathematical cancer modeling is currently increasing at a rate that is proportional to the rate of advancement in experimental oncology. However, to date, the mathematical models constructed in studies vary according to the type of cancer. For example, Sottoriva et al. proposed the “Big Bang model” for colon cancer (9), whereas Vermeulen et al. proposed a different model for colon cancer (10). Michor et al. established a model for imatinib therapy resistant cells in chronic myelogenous leukemia (11). Tomasetti and Vogelstein investigated the oncogenesis model in the context of stem cell divisions (12).

There are numerous reasons as to why different mathematical cancer models have been constructed. They include differences among different cancers, different experimental methods, and different methods of data analysis. Thus, there are still limitations in experimental oncology.

One method of overcoming these limitations is to establish a generalized mathematical model that can be applied to any type of cancer. Williams et al. successfully established such a model (13). This study approached model development from the perspective of a mathematician in order to establish a more generalized mathematical cancer model that can be applied to all types of cancer, thereby significantly enhancing the utility of mathematical cancer modeling and making it more useful to experimental oncologists.

To establish a fully generalized mathematical model of cancer, general cancer properties were mathematically interpreted before analysis of the data obtained from the experiments of a specific type of cancer. Then, actual data analyses were conducted to test whether the theory obtained in this work is valid. It was found that cancer growth follows the Yule process, and that a violation of the symmetry about the growth leads to tumor heterogeneity. Furthermore, Monte Carlo simulation was found to be an informative tool to analyze cancer growth.

## Methods

### Software used in this study

The R software was used to analyze data and simulate cancer cell growth. In addition, R was used to create all graphs except Fig. 2(a) and Fig. 3(a), which were drawn using Python, matplotlib.

### Data analysis

I re-analyzed the raw data published in (14) using unpublished data.

### Test of the distribution

As discussed in detail in the results section, the initial number of cancer cells, *n*_0_, and the growth rate of cancer cells, *λ*, are important parameters in theories. These parameters were calculated from Eqs. (7) and (8) using the raw data obtained above. Then, the distributions of *n*_0_ were indicated. To test if these distributions were Poisson distributions, the *χ*^2^ test and Fisher’s exact test were conducted with the null hypothesis that “the distribution of *n*_0_, which is calculated by Eq. (7), is equal to the Poisson distribution with *λ*, which is calculated by Eq. (8).” If *p* < 0.05, then this null hypothesis was set to be rejected.

### Design of the numerical solution and Monte Carlo simulation

This section describes the outline of the numerical solution and simulation conducted in this study. The detailed programs written for these calculations are available in the Supplementary Method section.

The numerical solution of *dN*(*t*) = *λN*(*t*)*dt* + *σdBt* is obtained by employing the Euler-Maruyama method: *n* partitions of the interval (0,*t*] are defined as (*t*_*i*−1_, *t_i_*], (*i* = 1,2,…, *n*), where Δ*t_i_*: *t*_*i*+1_ − *t_i_*. Then,

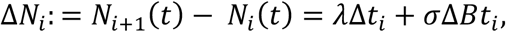

where Δ*Bt_i_* is a random number that follows a normal distribution with *mean* = 0 and 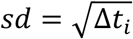. By solving the above equation, the numerical solution for *N*(*t*) is obtained: *N*(*t*) = *n*_0_ + ∑_*i*_Δ*N_i_*. This calculation is itself the Monte Carlo method (15). Based on the law of large numbers (15,16), repeating this calculation numerous times can reveal the population mean of cancer growth. This Monte Carlo calculation was repeated 100 times to simulate cancer growth; this is referred to as Monte Carlo simulations.

## Results

### Mathematical modeling of the growth of a cancerous tumor

First, the growth of cancer is assumed mathematically to follow the Poisson process with parameter *λ*. The Poisson process satisfies the following three conditions (16,17): 1) the increment for events occurring within the interval (*t, t* + *h*] is *λh;* 2) the process is time homogeneous, which implies that the distributions of the increments depend only on time and not on preceding scenarios; 3) each increment over contiguous time intervals is independent of the others.

In oncology, the growth of cancer appears autonomous because genetic and/or epigenetic abnormalities occur frequently in cell cycles or cell divisions, and the apoptosis of cancer cells occurs rarely, except in the cases of terminal cancer. Therefore, it is reasonable to assume that over a short period, the growth of cancer cells becomes almost constant, independent of preceding conditions, and occurs at random, i.e., it follows the Poisson process.

Let *N*(*t*) be the number of cancer cells at time *t*, and *P*[*N*(*t*) = *n*] (abbreviated as *p_n_*(*t*)) be the probability that *N*(*t*) = *n* at time *t*. The change in the number of cancer cells over a short interval (*t, t* + *h*] is calculated under the above conditions.

For an interval (*t, t* + *h*], probability *P*[*N*(*t* + *h*) = *n*] can be expressed as the sum of three states. Two of these states are as follows: 1) the probability of no cell division occurring in the interval (*t, t* + *h*] under the condition *N*(*t*) = *n* and 2) the probability of one cell division occurring in the interval (*t, t* + *h*] under the condition *N*(*t*) = *n*−1. The remaining probabilities include one in which the number of cells increases by two or more as a result of one instance of cell division or cell death. However, this state is rarely observed in oncology because of the assumption described above; thus, the other probabilities are close to zero.

As these three states are independent of one another and the increase in the number of cells at time interval *h* under the condition *N*(*t*) = *n* is *nλh*, the probabilities for the interval (*t, t* + *h*] can be given by

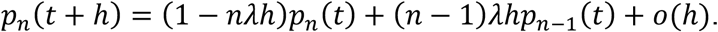

Rearranging this equation and taking the limit *h* → 0 yields the following:

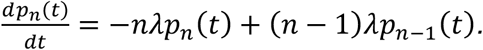

Let *n*_0_ = *N*(0); then, the initial condition can be defined as

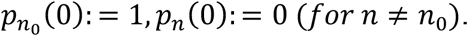

It is known that this differential equation has the following solution:

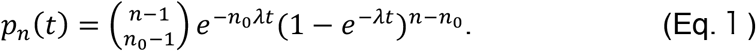

This is a negative binomial distribution; thus, the expectation and variance are already known as

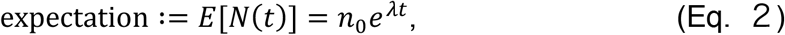

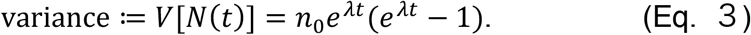

These results represent the Yule process, or the pure birth process, which is a model of probability that has been extensively studied in mathematics and applied to various other areas including physics and biology. Interested readers can refer to (16) for details. The negative binomial distributions of cancer cells are shown in Figs. 1(a) and (b).

**Fig. 1.**
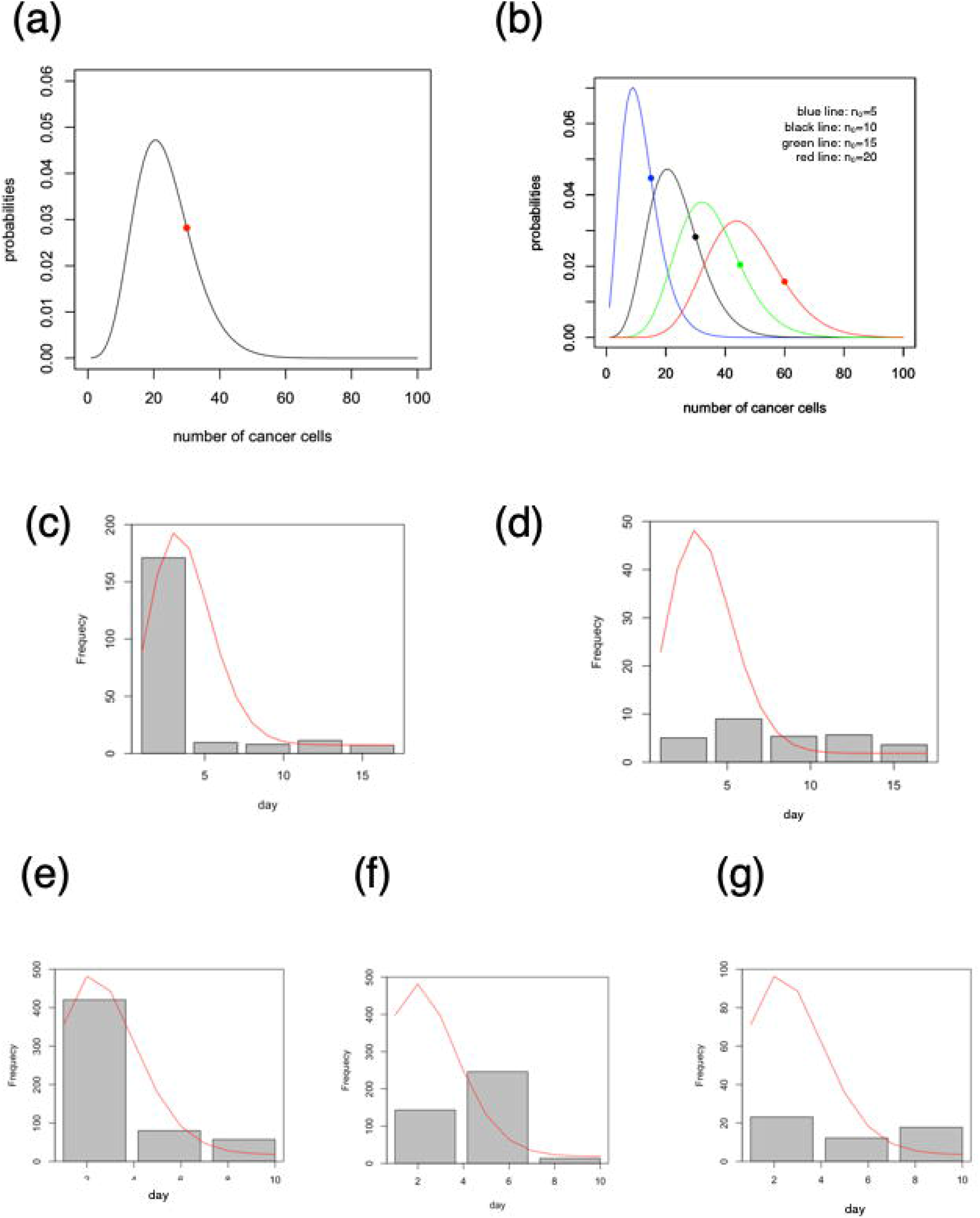
Distribution of a cancer. (a) Negative binominal distribution of cancer cells with *n*_0_ = 10. The red point indicates the expected value. The number of cancer cells at the peak of this distribution are always below that expected, which is characteristic of negative binomial distribution. (b) Change in distribution with variable *n*_0_, whose value ranges from 5 to 20. The dot on each graph indicates the expected value. It was found that the height of the peak decreases as the value of *n*_0_ increases From (c) to (g) The distribution of *n*_0_ (bar graph) and Poisson distribution with *λ* (red line plot); both parameters were calculated using Eqs. (7) and (8), respectively. (c) Graphs of H226 CV cell lines. This bar graph distribution is obviously different from normal distribution, but seems to be equal to the Poisson distribution (red line). (d) Graphs of H226 sh-CD24 cell lines. (e) Graphs of JMM CV cell lines. (f) Graphs of JMM sh-CD24 cell lines. (g) Graphs of JMM-shCD26 cell lines.

### Solving the stochastic differential equation

Eq. (2) expresses that the expected number of cancer cells at time *t* is *n*_0_*e*^*λt*^ when the number of cancer cells at time 0 is *n*_0_. This demonstrates that cancer is expected to grow exponentially. Differentiating Eq. (2) yields the following equation:

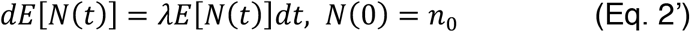

Even though I have assumed above that the growth of cancer cells is random and the distributions of their increments depend only on the time interval, Eq. (2’) indicates that the randomness of the growth is zero in the short period if we take the average of the number of cancer cells. Thus, I introduce the stochastic process, whose average is zero and whose variance depends only on time intervals. The Brownian motion function is introduced to the growth formula of cancer cells because it is well known that the average of this function is zero and the variance of this function depends only on the time interval (16). In brief, it is natural to introduce the Brownian motion function to the growth formula of cancer cells based on the above assumption.

Supposing that the randomness of the growth rate of cancer follows the Brownian motion function, the change in the number of cancer cells can be intuitively expressed as

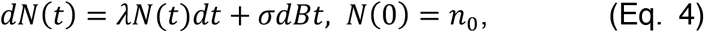

where *Bt* denotes the Brownian motion function and *σ* is a deterministic function, which is assumed to be independent of time for the sake of simplicity in this study. A detailed introduction to this model is provided in (17). In the current study, this model has been applied to the field of oncology.

Eq. (4) is known to have a solution (17). Using Ito calculus, the solution can be expressed as

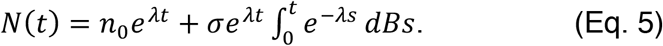

This is a form of the Langevin equation and an example of the Ornstein–Uhlenbeck process (17). The average of the integral in the second term on the right-hand side of Eq. (5) is zero, and its variance is 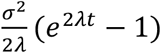 (17). This variance must be equivalent to Eq. (3); thus,

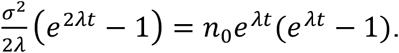

This equation is solved for *σ*, and 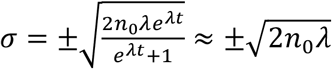.

For the sake of simplicity, it has been assumed above that *σ* is time independent; thus, this equation does not contradict the above assumption.

Therefore, Eq. (5) can be expressed as

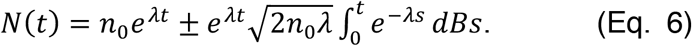

For simplicity,

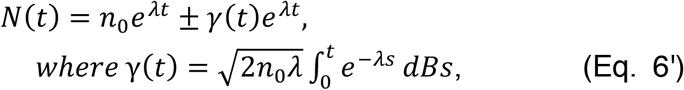

where *γ*(*t*) is the function following Brownian motion, *n*_0_ is the initial number of cancer cells, and *λ* is the growth rate.

### Application of the model to the in vitro data

In general, if we perform appropriate experiments or the observations of cancer patients’ status, then we obtain the mean and variance of cancer growth. 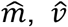, and 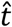 are defined to be the actual values of mean, variance, and time, respectively. Note that these three symbols with the hat mark are not algebraic variables but the actual experimental data obtained. Then, Eqs. (2) and (3) can be written as

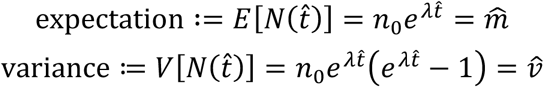

As these two equations are simultaneous linear equations about *n*_0_ and *λ*, these two variables can be solved as follows:

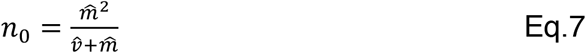

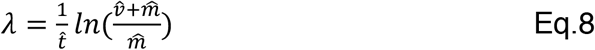

Eqs. (7) and (8) indicate that we can calculate *n*_0_ and *λ* from actual data if we perform appropriate experiments or observations.

Now, the theories discussed above are tested by applying them to the actual in vitro data obtained from cancer growth experiments. It was reported that CD24 and CD26 are suggested as the markers of cancer stem cells in malignant pleural mesothelioma (14). In this report, we established the knock down cells of CD24 or CD26 using short hairpin RNA (they are referred to as sh-CD24 cell lines and sh-CD26 cell lines, and control cell lines are referred to as CV cell lines). We performed the growth test of two different mesothelioma cell lines, i.e., JMM cell lines and H226 cell lines, and compared them with knock down of CD24 or CD26 (14). I re-analyzed these results using unpublished data. The reanalysis data and the calculations of *n*_0_ and *λ* are listed in Table 1.

**Table 1.** Calculation data of the growth of five cell lines. The means and variances of the five cell lines are cited in (14) with unpublished data. The values of *n*_0_ and *λ* of each cell line are calculated using Eqs. (7) and (8).

|  | DAY1 | DAY4 | DAY7 | DAY10 | DAY14 | DAY17 |
| --- | --- | --- | --- | --- | --- | --- |
| MEAN<br>(UNIT: $\times 10^5$) | 1.00 | 3.47 | 9.79 | 29.52 | 146.94 | 299.77 |
| VARIANCE<br>(UNIT: $\times 10^5$) | 0 | $3.56 \times 10^3$ | $4.17 \times 10^5$ | $5.27 \times 10^6$ | $1.26 \times 10^8$ | $4.47 \times 10^8$ |
| $n_0$ | N/A | 171.01 | 9.68 | 8.02 | 11.46 | 7.27 |
| $\lambda$ | N/A | 2.42 | 3.70 | 4.14 | 3.46 | 4.85 |

**(b) Growth data of H226 sh-CD24 cell lines.**
|  | DAY1 | DAY4 | DAY7 | DAY10 | DAY14 | DAY17 |
| --- | --- | --- | --- | --- | --- | --- |
| MEAN (UNIT: $\times 10^5$) | 1.00 | 1.33 | 3.98 | 12.85 | 35.61 | 62.27 |
| VARIANCE<br>(UNIT: $\times 10^5$) | 0 | $2.20 \times 10^3$ | $7.99 \times 10^4$ | $1.54 \times 10^6$ | $1.07 \times 10^7$ | $3.66 \times 10^7$ |
| $n_0$ | N/A | 5.06 | 8.99 | 5.38 | 5.68 | 3.62 |
| $\lambda$ | N/A | 2.93 | 3.43 | 4.00 | 3.23 | 4.55 |

**(c) Growth data of JMM CV cell lines.**
|  | DAY1 | DAY4 | DAY7 | DAY10 |
| --- | --- | --- | --- | --- |
| MEAN<br>(UNIT: $\times 10^5$) | 1.00 | 8.93 | 23.46 | 79.63 |
| VARIANCE<br>(UNIT: $\times 10^5$) | 0 | $1.49 \times 10^2$ | $2.79 \times 10^3$ | $5.79 \times 10^5$ |
| $n_0$ | N/A | 420.48 | 79.90 | 57.23 |
| $\lambda$ | N/A | 1.75 | 2.50 | 3.06 |

**(d) Growth data of JMM sh-CD24 cell lines.**
|  | DAY1 | DAY4 | DAY7 | DAY10 |
| --- | --- | --- | --- | --- |
| MEAN<br>(UNIT: $\times 10^5$) | 1.00 | 7.37 | 13.47 | 29.18 |
| VARIANCE<br>(UNIT: *10<sup>5</sup>) | 0 | 28.22 | 433.10 | 1.89*10<sup>4</sup> |
| $n_0$ | N/A | 143.30 | 246.02 | 13.35 |
| $\lambda$ | N/A | 2.03 | 1.84 | 3.12 |

**Table 1 Calculation data of the growth of five cell lines.**
|  | DAY1 | DAY4 | DAY7 | DAY10 |
| --- | --- | --- | --- | --- |
| MEAN<br>(UNIT: *10<sup>5</sup>) | 1.00 | 4.40 | 10.45 | 24.04 |
| VARIANCE<br>(UNIT: *10<sup>5</sup>) | 0 | 500.00 | 3.49*10<sup>3</sup> | 1.39*10<sup>4</sup> |
| $n_0$ | N/A | 23.10 | 12.23 | 17.73 |
| $\lambda$ | N/A | 2.43 | 2.84 | 2.98 |

Interestingly enough, even though the growth tests were started by seeding 10^5^ cells in all cell lines, the calculations show that *n*_0_ is at most approximately 500 at each day and below ten in certain cases (Table 1). Additionally, the *n*_0_ at each day decreases considerably as the day passes. These distributions are shown in Figs. 1(c) to (g). The Poisson distributions with parameter *λ*, which is calculated by Eq. (8), are also shown in Figs. 1(c) to (g) by red lines. To test whether the distributions of *n*_0_ at each day are Poisson distributions, *χ*^2^ tests were conducted with the null hypothesis that “the distribution of *n*_0_, which is calculated by Eq. (7), is equal to the Poisson distribution with *λ*, which is calculated by Eq. (8)” (16). The results of this test reveal that two H266 cell lines (CV cell lines and sh-CD24 cell lines) are calculated as *p* = 0.22 and three JMM cell lines (CV cell lines, sh-CD24 cell lines, and sh-CD26 cell lines) are calculated as *p* = 0.20. Thus, the null hypothesis cannot be rejected. However, as R warned that *χ*^2^ tests might be incorrect in these situations, Fisher’s exact tests were additionally conducted for the same samples under the same conditions. The results of these tests are *p* = 1 in all five cell lines. The two tests prove that the distribution of *n*_0_ is almost surely the Poisson distribution. Thus, it is concluded that the growths of these five cell lines follow the Poisson process and that the abovementioned theories can be applied to these cell lines.

Based on these results, I exploited the data of the H226 and JMM cell lines to solve Eq. (6) numerically. The results are plotted in Fig. 2 and Fig. 3, while the actual data obtained from (14) are shown in Fig. 2(a), and Fig. 3(a). Monte Carlo simulations reveal that the growth of sh-CD24 cell lines and sh-CD26 cell lines is evidently slower than that of CV cell lines, which is in agreement with actual data (Figs. 2 and 3). The results of the Monte Carlo simulations reinforce the former report and demonstrate that the theories obtained from the above discussion can be verified through in vitro experiments. These theories are suggested to be valid in the five cell lines.

**Fig. 2.**
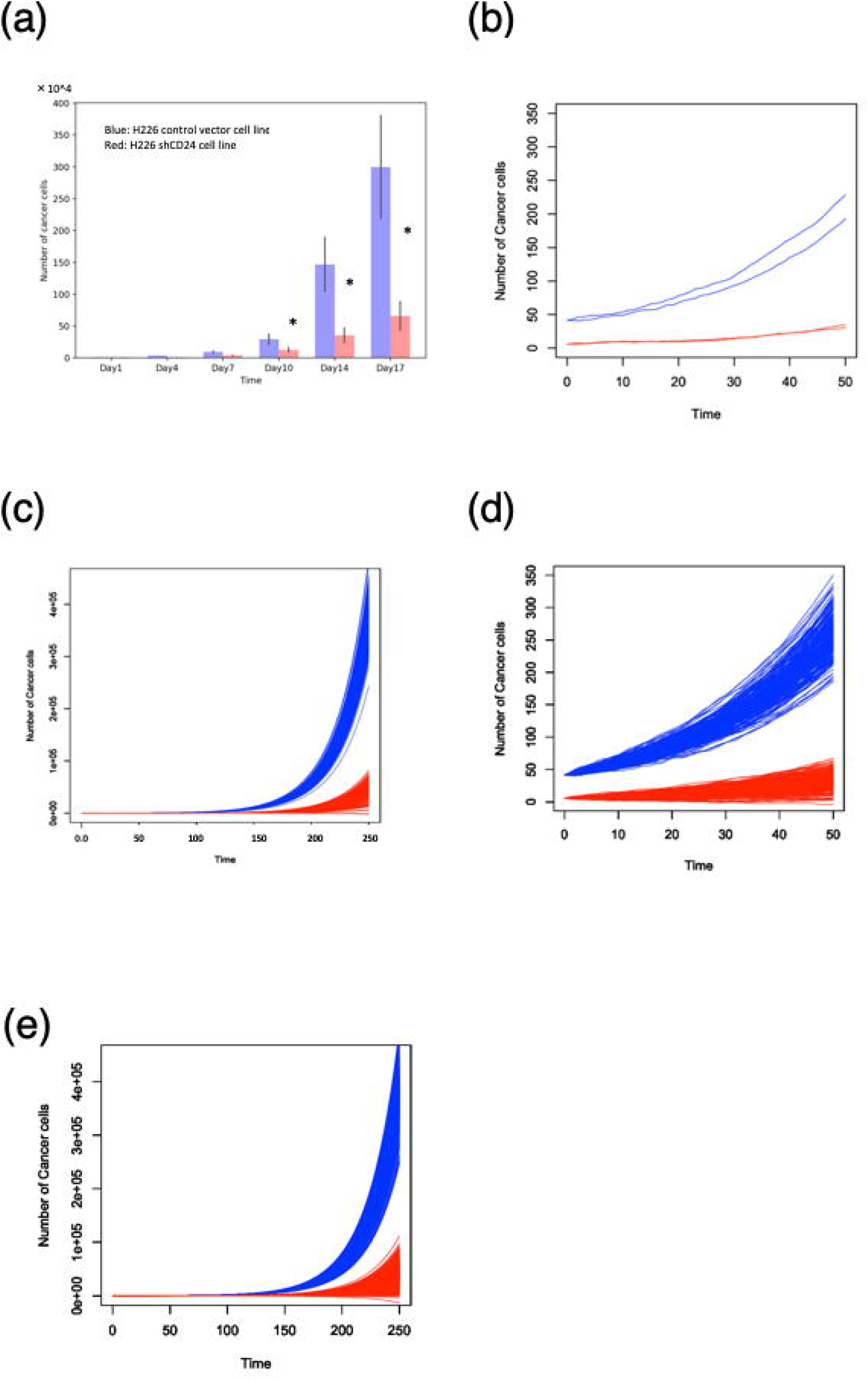
Comparison of cancer cell (H226) growth between actual data obtained from the experiment and simulation experiments. (a) Comparison of growth between H226 CV cell lines (blue bar graph) and H226 sh-CD24 cell lines (red bar graph). * indicates the significant difference between the CV cell lines and sh-CD24 cell lines. This graph is basically equal to the one published in (14) (b) Numerical solution of the two cancer cell lines by Eq. (6). The time interval is set to (0,50]. The blue plots denote the solutions of the H226 CV cell lines, while the red plots denote the solutions of the H226 sh-CD24 cell lines. Both pathways are not smooth but have a zigzag pattern. This is the characteristic of the stochastic differential equations (17). (c) Monte Carlo simulation of H226 cells lines. the simulation calculations were performed 100 times. The time interval was set to (0,250]. The growth of sh-CD24 cell lines is obviously slower than that of CV cell lines. The notation of the Y-axis value, i.e., 1e+05, means 1.0*10^5^ (this notation is usually used in R and will be used later) (d) Magnification of (c). The time interval was set to (0,50]. (e) Monte Carlo simulation of H226 cell lines. The results of performing the calculation 500 times are shown. This graph is similar to that of shown in (c). Thus, the graphs (c) and (e) suggest that performing the calculation 100 times is sufficient to simulate the growth of a cancer. Some pathways of red plots are found to be nearly zero.

In conclusion, if the mean and variance of cancer growth are obtained through appropriate experiments, *n*_0_ and *λ* can be calculated by Eqs. (7) and (8). The distribution of *n*_0_ calculated by Eq. (7) is demonstrated to be almost surely the Poisson distribution with *λ* calculated by Eq. (8). Thus, *N*(*t*) can be obtained by numerically solving Eq. (6). Monte Carlo simulations are suggested to be an informative tool to analyze the growth of a cancer.

### Violation of symmetry by cancer clones leads to cancer heterogeneity

The interpretations of Eq. (1) and Eq. (6) are continued in this section. First, Eq. (6) apparently consists of two equations. Thus, Eq. (6) indicates that cancer growth follows two different formulae. *N*(*t*) = *n*_0_*e^λt^* + *γ*(*t*)*e^λt^* is defined as the Ryu pathway and *N*(*t*) = *n*_0_*e^λt^* − *γ*(*t*)*e^λt^* as the Ken pathway. Eq. (6) is numerically solved with *n*_0_ = 1 to observe how cancer occurs from a single cell. Based on the former section, it is reasonable to set the growth rate as *λ* = 2.5. Figs. 4(a) to (d) show the results of the simulation of Eq. (6) with *n*_0_ = 1. These figures show that two pathways emerge from a “single” cancer cell. To avoid misunderstanding, in this study, symmetric clones have been defined as the clones with identical genetic expressions and that occur in epigenetic scenarios in an environment. Figs. 4(a) and (b) show that the different growth patterns that follow Eq. (6) exist from a single clone. Fig. 4(a) shows that the clones proliferate competitively. A few competitive proliferations are found in Fig. 2(c) and Fig. 3(c). If the clones that follow a pathway (e.g., Ryu) can compete with the other clones (e.g., Ken), then the majority of cancer cells consist of Ryu clones and Ken clones form a minor population (Fig. 4(c)). Thus, Eq. (6) states that a clone can produce asymmetric cell populations.

**Fig. 3.**
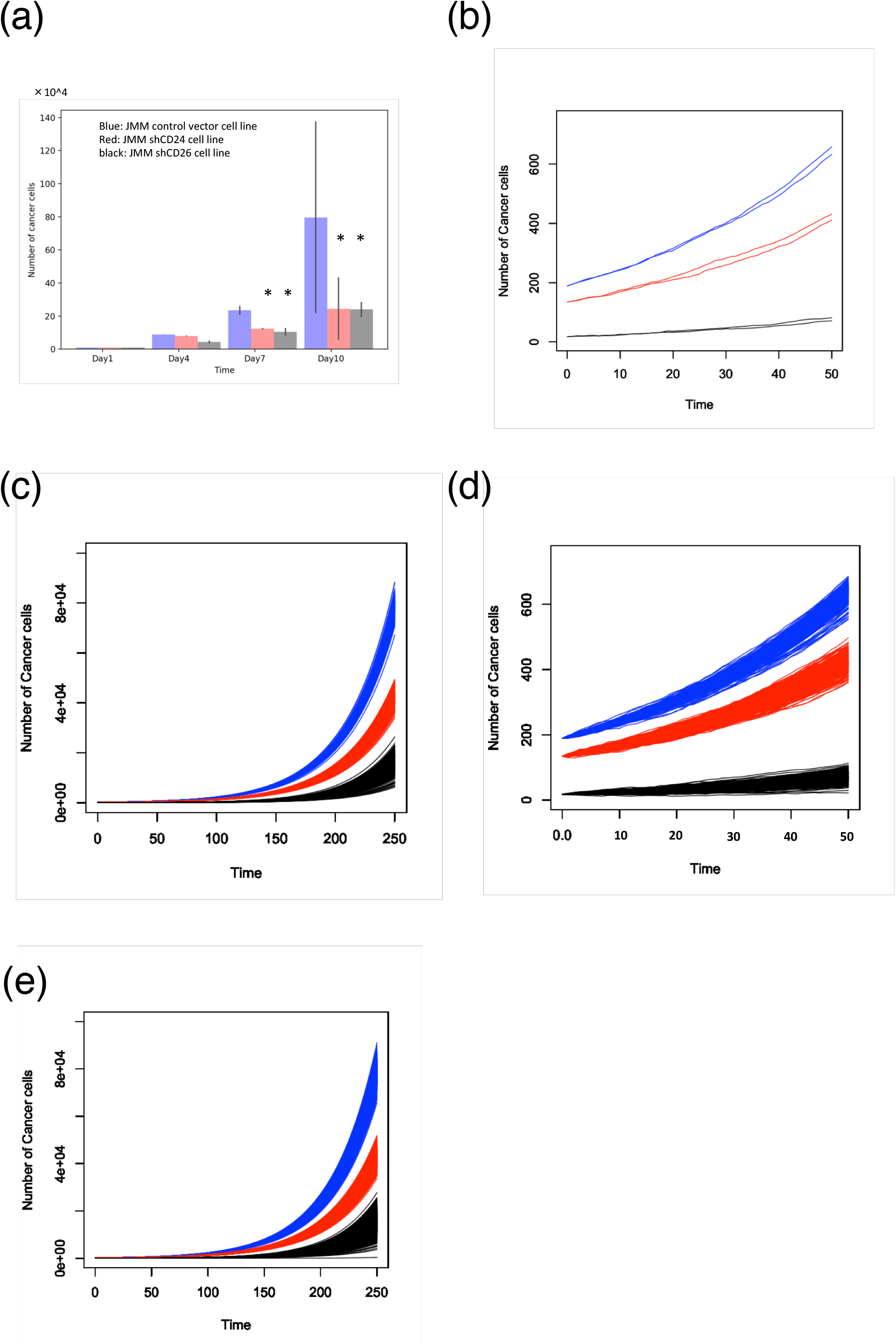
Another comparison of cancer cell (JMM) growth between actual data and those of simulation experiments. (a) Comparison of the growth between JMM CV cell lines (blue bar graph) and H226 sh-CD24 cell lines (red bar graph), and JMM sh-CD26 cell lines (black bar graph). * indicates the significant difference between the CV cell lines and sh-CD24 or sh-CD26 cell lines. This graph is identical to the one published in (14) as well. (b) Numerical solution of the three cancer cell lines. The blue plots denote the solutions of the JMM CV cell lines, the red lines denote the solution of the JMM sh-CD24 cell lines, and the black cell lines denote the solution of the JMM sh-CD26 cell lines. (c) Monte Carlo simulations of three cell lines. The numerical solutions are repeated 100 times. The time interval was set to (0,250]. The growth of the sh-CD26 cell lines is obviously slower than those of the other two lines, and the growth of the sh-CD24 cell lines is obviously slower than that of the CV cell lines. (d) Magnifications of (c). The time interval was set to (0,50]. (e) Monte Carlo simulation of H226 cell lines. The results of performing the calculation 500 times are shown. This graph is similar to that shown in (c). Thus, the graphs (c) and (e) suggest that performing the calculation 100 times is sufficient to simulate the growth of a cancer as well as that of H226 cell lines. Some pathways of black plots are found to be nearly zero.

**Fig. 4.**
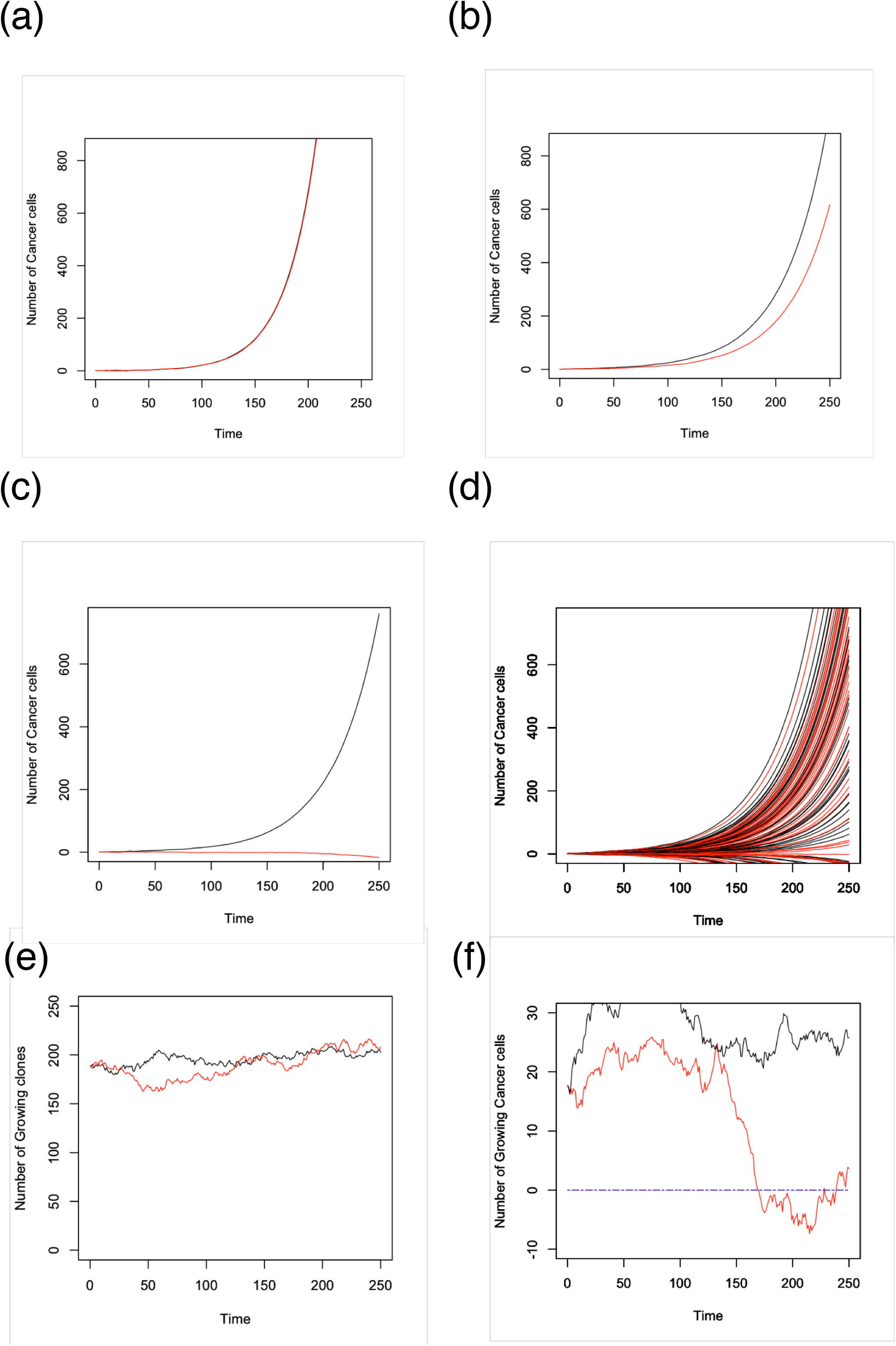
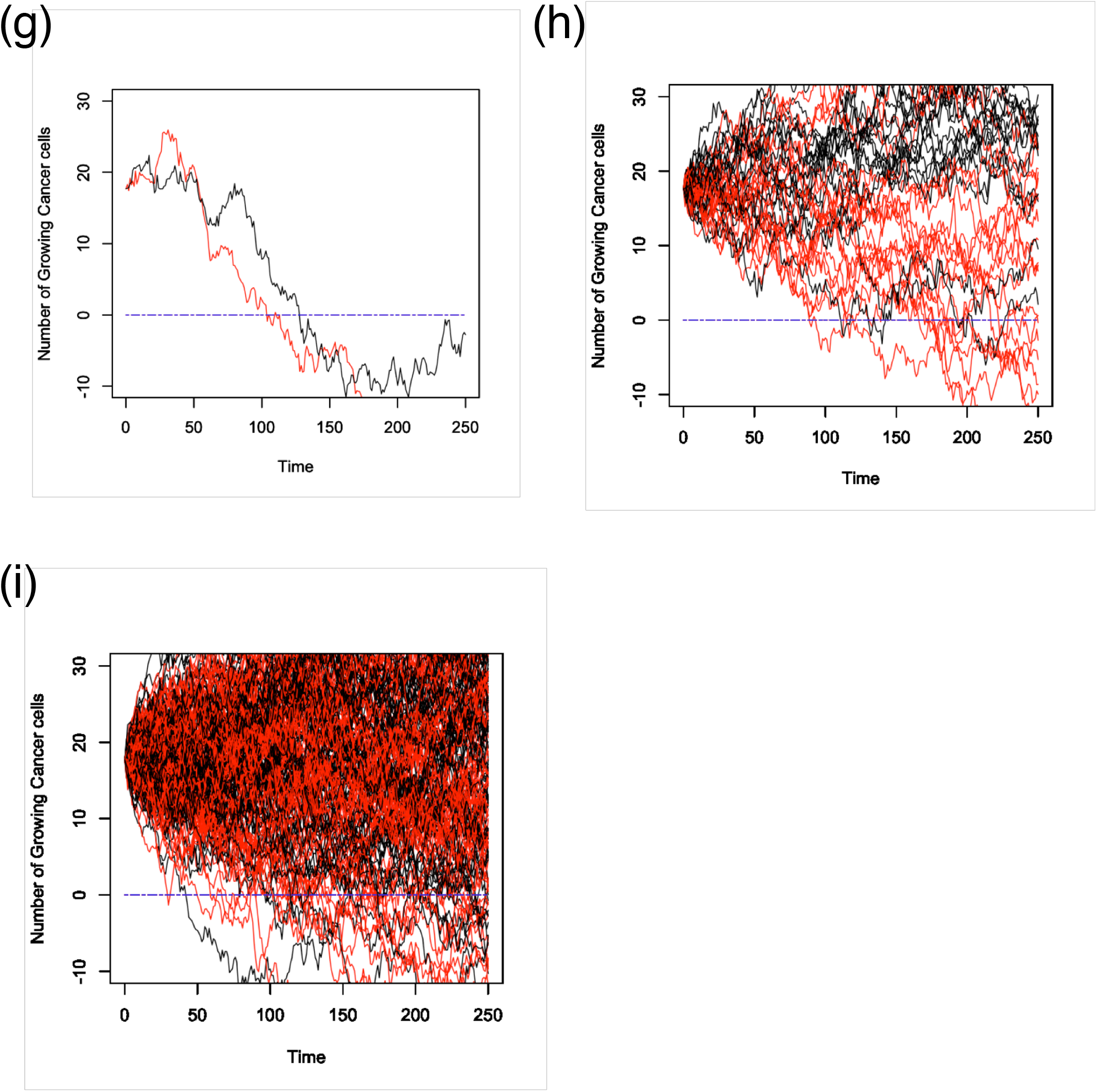
Simulation experiments that show the symmetry violation of cancer cells. Equation (6) is numerically solved for *n*_0_ = 1 and *λ* = 2.5. The black line indicates the Ryu pathway, and the red line indicates the Ken pathway. (a) Growth simulations of both the pathways. This graph shows that both the pathways grow competitively. (b) The simulation result shows that the numbers of Ryu pathways are always greater than those of Ken pathway. (c) The graph that denotes the growth of the Ken pathway is quiescent whereas that of the Ryu pathway grows normally. (d) The simulation experiments were repeated 50 times. Notice that there are some pathways in which the cell numbers go below zero. (e) Numeric solution of the number of growing clones *N_g_*(*t*) in the JMM CV cell lines. The two plots consist of the Ryu pathway and Ken pathway. These graphs apparently show that the mean of the growing clone is *n*_0_. (f) Numerical solution of the number of growing clones *N_g_*(*t*) in the JMM sh-CD26 cell lines. The blue dashed line indicates the zero line. It can be observed that the red line goes below zero once and then goes above zero again. The red line decreases drastically during the short time it is observed at time [130,160] as well. (g) Another solution of *N_g_*(*t*) of the JMM sh-CD26 cell lines. Both lines decrease globally over time, and the number of growing clones is less than zero halfway through the observation period.

Fig. 4(c) shows the simulation data for the case in which the number of cancer cells that followed the Ken pathway could not proliferate for some reason; this implies that Ken clones’ proliferations are quiescent. If the cancer cells that did not exhibit proliferation were allowed to proliferate, Eq. (6) would repeatedly produce asymmetric clones. Fig. 4(d) shows the results of Monte Carlo simulations with calculations performed 50 times. The figure shows that a few pathways are below zero. This suggests that cancer growth has the potential to stop autonomously. The same situations can be observed to a minor extent in Figs. 2(c) and 3(e).

Let us now focus on the interpretation of Eq. (1). Considering *q* = *e*^−*λt*^, Eq. (1) is rewritten as follows:

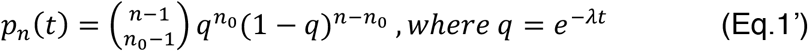

Mathematically, Eq. (1’) is typically interpreted as follows: there is a trial with two potentials referred to as “success” and “failure,” where the probability of success is *q*. Then, *p_n_*(*t*) expresses the probability of the occurrence of *n*_0_ successes in *n* trials (16). Based on this concept, I interpret Eq. (1’) as follows: cancer clones always select “success,” that is, to proliferate, or “failure,” that is, not to proliferate, every time. The probability of a clone that proliferates at time *t*, which is named as a growing clone, is *q* = *e*^−*λt*^. Then, *p_n_*(*t*) expresses the probability that *n*_0_ clones out of *n* cancer cells “succeed” in proliferating. Probability *q* indicates that the number of growing clones decreases considerably as time passes. This finding is in agreement with the distribution of *n*_0_ (Figs. 1(c) to (g)). If there exists a clone whose probability *q* is nearly zero, then such a clone appears to be quiescent. However, such a clone has the potential to proliferate because *q* is not exactly zero at any time.

Let *N_g_*(*t*) be the number of growing clones at time *t. N_g_*(*t*) is defined as

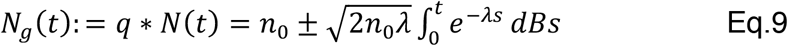

Eq. (9) suggests that *N_g_*(*t*) ≤ 0 could be possible depending on parameters. *N_g_*(*t*) ≤ 0 implies that a cancer cannot grow in a short time interval. At the time period in which *N_g_*(*t*) ≤ 0, cancer is globally quiescent. As the average of the Brownian motion function is zero,

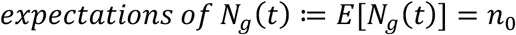

holds. This equation justifies that *n*_0_ is a one of the parameters of cancer. According to Eq. (1’), cancer globally always endeavors to make *n*_0_ clones proliferate at any time.

To investigate if the theories discussed in this section can be applied to actual data, Eq. (9) is numerically solved using JMM CV cell lines and JMM sh-CD26 cell lines. While the *n*_0_ of JMM CV is the largest among the five cell lines, the *n*_0_ of JMM sh-CD26 cell lines is the smallest (Table 1). This is the reason for using the two cell lines in this numerical solution. The calculation result for JMM CV cell lines shows that *N_g_*(*t*) is always approximately 150 during the time interval (0, 250] (Fig. 4(e)). Therefore, growing clones exist in JMM CV cell lines at almost every time in (0, 250]. This result suggests that it is justified that *n*_0_ is the parameter of cancer. In contrast, in JMM sh-CD26 cell lines, a few clones are calculated as *N_g_*(*t*) ≤ 0 (Figs. 4(f) to (i)). Moreover, a few growing clones are calculated once *N_g_*(*t*) ≤ 0 and after considerable time, *N_g_*(*t*) > 0 has the potential to re-occur (Fig. 4(f)). This result suggests that cancer clones always have the potential to proliferate at any time even if *N_g_*(*t*) ≤ 0. This result reinforces the result shown in Fig. 4(d) and its theory.

Considering the above results, Eq. (6) is shown to describe the violation of symmetry of cancer cells that leads to cancer heterogeneity. Eq. (1) also states that cancer heterogeneity globally includes the fact that cancer is quiescent in a few cases. Eq. (9) describes the number of growing clones. Considered together, Eq. (1), Eq. (6), and Eq. (9) are the equations that describe cancer heterogeneity.

## Discussion

In this study, a mathematical model was developed by implementing a stochastic process based on the mathematical interpretation of generalized cancer properties. It was shown that cancer growth follows the Yule process. Thus, when there are *n*_0_ cancer cells at time 0, the expectation and variance of cancer growth at time *t* can be determined via Eqs. (2) and (3).

First, I assumed that the growth of the cancer cells occurred at random in a short time period. Eq. (2’) showed that the randomness was zero when one calculated the mean of cancer growth. Therefore, it was reasonable to suppose such randomness followed a Brownian motion function, and Eqs. (4) to (8) were obtained. If we perform appropriate experiments or the observations of cancer patients’ status, then we can obtain the mean and variance of cancer growth. In this study, I found the formulae of *n*_0_ and *λ* to calculate from the mean and variance obtained appropriate experiments (Eqs. (7) and (8)). I proved that the distribution of *n*_0_, which was calculated by Eq. (7), was almost surely the Poisson distribution with parameter *λ*, which was calculated by Eq. (8) using the experimental data. Based on this proof, Eq. (6) was solved numerically. The results of the numerical solution of Eq. (6) and its Monte Carlo simulations reinforced the former report and demonstrated that the theories obtained from above discussion can be verified via in vitro experiments. Further, Monte Carlo simulations were suggested to be an informative tool to analyze the growth of a cancer. As my assumption is simple and can be adapted to the all kinds of cancers, Eq. (6) and its Monte Carlo simulations can be adapted to all kinds of cancer and can provide informative results. However, more experimental investigations are necessary to apply the equation to all type of cancers.

In physics, randomness is referred to as additive noise (17). It is natural to consider that the source of this noise (randomness) generally consists of two factors in oncology: an intrinsic factor and an extrinsic factor. Elowitz et al. stated that E.coli could yield different responses to these two factors (18). It would be interesting to investigate whether these results are applicable to cancer cells. A major intrinsic factor in a cell is the instability of a genome. Williams et al. considered the instability of a genome to be pink noise, which is common in nature (13). Nowell discussed cancer heterogeneity in the context of genome instability (19). In addition, Abdallah et al. recently reported that the distribution of cancer cells with an unstable genome is characteristic, whereas cells with a stable genome have a relatively normal distribution (20). Even though they only stated that the distribution is characteristic, I consider that the distribution appears to be the negative binominal distribution shown in Figs. 1(a) and (b). The results on intrinsic factors continue to accumulate; however, extrinsic factors are largely neglected. To take these facts into account, it is necessary to examine whether the randomness in cancer growth is influenced by intrinsic factors, extrinsic factors, or both.

It is found that Eq. (6) can produce asymmetric clones (Figs. 2(b) and (c), Figs. 3(b) and (c), and Fig. 4(a) to (c)). A simulation experiment shows that asymmetric clones can be produced by a single cancer cell, which is followed by Eq. (6) (Figs. 4(a) and (b)). Asymmetric clones typically appear to proliferate competitively (Fig. 2(b), Fig. 3(b), and Fig. 4(a) and (b)). Such a locally competitive asymmetric proliferation will cause cancer heterogeneity globally.

Eq. (1’) can be interpreted as follows: cancer clones always select “success,” that is, to proliferate, or “failure,” that is, not to proliferate, every time. The probability of a growing clone is *q* = *e^−λt^*. Then, *p_n_*(*t*) expresses the probability that *n*_0_ clones out of *n* cancer cells “succeed” in proliferating. The number of growing clones, *N_g_*(*t*), can be formulated as Eq. (9). Fig. 4 demonstrates that the events of *N_g_*(*t*) ≤ 0 are not rare events depending on two parameters. *N_g_*(*t*) ≤ 0 implies that growing clones never exist in the particular time period, that is, cancer is globally quiescent at this time. The probability of *q* is interesting because it indicates that success probability decreases exponentially as time passes. In accordance with this, the Poisson distribution expresses that *n*_0_ decreases considerably as time passes. Considered together, the cancer clones that can proliferate are suggested to decrease considerably as time passes. Figs. 1(a) and (b) show that the numbers of cancer cells at the peak of the distribution are always below expectation, which is the property of negative binomial distributions (16). Thus, Eq. (1) implies that the majority of cancer clones, which include quiescent clones, grow slower than expected. One of the reasons for this might be asymmetric divisions because locally competitive proliferations make cancer growth slower. Considered together, Eqs. (1), (6), and (7) represent tumor heterogeneity, which is caused by the violation of symmetry.

The Yule process was originally established by Yule to mathematically formulate Darwin’s evolution theory (16,21). Subsequently, this process became an important topic of study in mathematics and was applied in several fields of science (16). In recent years, McGranahan and Swanton interpreted that tumor evolution within a cancer patient might follow the pattern of the evolution of species according to Darwin’s evolution theory (22). As the cancer model established in this study follows the Yule process (such as Eq. (1)) and it is developed further into Eq. (6), in agreement with McGranahan and Swanton, it may be suggested that cancer evolution follows Darwin’s evolution theory. However, further experimental investigations are necessary to substantiate this hypothesis.

In conclusion, in this study, a generalized mathematical model of cancer was developed based on the general properties of cancer cells and an equation was derived to represent the violation of symmetry of cancer clone cells that leads to cancer heterogeneity. Re-analysis of the experimental data validated the theories. Monte Carlo simulation was found to be an informative tool for analyzing cancer growths. These findings are expected to provide a deeper understanding of the mechanisms of cancerous tumor growth. Additionally, with further experimentation, the simulation of the growth of cancer cells presented here has the potential to reveal each cancer’s own specific features or common properties among the different kinds of cancers.

## Supporting information

supplemental data using to creat a the Table etc.

## Acknowledgment

The author is grateful to his family and friends for support. Masayuki Yamamoto, a friend who is a computer programmer, offered valuable advice and wrote the beautiful code for the growth simulation. The author would also like to thank Editage (*www.editage.jp* for English language editing.

