## supplemental methods for "Generalized mathematical model of cancer heterogeneity"

**Motohiko Naito**

**Affiliations**: Ronin (Not affiliated with any institute)

**Correspondence:**

**Supplementary Methods**

**Code of growth simulation.**

Note that the code mentioned below has a random number in it. Therefore, even if you run this code with no arrangement, it is not possible to obtain the exact figures presented in this paper.

The codes show in Figs 2 and 3 are similar. Thus, only the code shown in Fig. 3 is presented in this manuscript (if you want the data of Fig 2 , then you only exchange the value of lam, sigma, and n (appeared first time))

**Fig 3 code**

**#Generating the function that plots the Ryu pathway of the JMM CV cell lines.**

\###
\# jmmryucv ()
\###

jmmryucv <-function(){

#fixing the time and parameters

dt <- 0.01 # time interval

TT <-250 # end time. TT is 50 in some figures.

lam <-2.44 # growth rate: lambda, one of the parameters of this equation

sigma <-18.73*sqrt(dt) # sigma is the coefficient of the Brownian motion function.

n <-188.87 #substituting the initial number of cancer cells, another parameter of this equation

#making the list of results

nn<-n

tt <- 0.0

#main body of this calculation

for(i in 1:TT){

#calculating the increment. The second term says that “Create the one random number that follows normal distribution and multiply it by sigma.”

dn <-(lam*n)*dt+sigma*rnorm(1)

n <-dn+n

\# if variable i is integral, the latest results are added to the list nn and tt

if(i %% 1==0){

nn<-rbind(nn,n)

tt<-rbind(tt,i);

}

}

#plot the results of the calculations

matplot(tt,nn,type="l",ylab="Number of Cancer cells", xlab="Time",col="blue”, ylim=c(0,10000))

}

**#Generating the function that plots the Ken pathway of JMM CV cell lines**

\###
\# jmmkencv ()
\###

jmmkencv <-function(){

dt <- 0.01

TT <-250

lam <-2.44

sigma <-18.73*sqrt(dt)

n <-188.87

nn<-n

tt <- 0.0

for(i in 1:TT){

dn <-(lam*n)*dt-sigma*rnorm(1)

n <-dn+n

if(i %% 1==0){

nn<-rbind(nn,n)

tt<-rbind(tt,i);

}

}

matplot(tt,nn,type="l",ylab="Number of Cancer cells", xlab="Time",col="blue",ylim=c(0,1e+05))

}

＃**Generating the function that shows the two pathways in one sheet**

\###
\# jmmcvplot ()
\###

jmmcvplot <-function(){

jmmkencv()

par(new=TRUE)

jmmryucv()

}

\# **Calculation codes of the JMM sh-CD24 cell lines, which is basically the same as that of the JMM CV cell lines.**

\###

\# jmmsd24ryu()

\###

jmmsd24ryu <-function(){

dt <- 0.01

TT <-250

lam <-2.33

sigma <-21.11*sqrt(dt)

n <-134.22

nn<-n

tt <- 0.0

for(i in 1:TT){

dn <-(lam*n)*dt-sigma*rnorm(1)

n <-dn+n

if(i %% 1==0){

nn<-rbind(nn,n)

tt<-rbind(tt,i);

}

}

matplot(tt,nn,type="l",ylab="Number of Cancer cells", xlab="Time",col="red",ylim=c(0,1e+05))

}

\###

\# jmmsd24ken()

\###

jmmsd24ken <-function(){

dt <- 0.01

TT <-250

lam <-2.33

sigma <-21.11*sqrt(dt)

n <-134.22

nn<-n

tt <- 0.0

for(i in 1:TT){

dn <-(lam*n)*dt+sigma*rnorm(1)

n <-dn+n

if(i %% 1==0){

nn<-rbind(nn,n)

tt<-rbind(tt,i);

}

}

matplot(tt,nn,type="l",ylab="Number of Cancer cells", xlab="Time",col="red",ylim=c(0,1e+05))

}

\###

\# jmmsd24plot()

\###

jmmcd24plot <-function(){

jmmsd24ken()

par(new=TRUE)

jmmsd24ryu()

}

＃**Calculation codes of the JMM sh-CD26 cell lines**

\###

\# jmmsd26ryu()

\###

jmmcd26ryu <-function(){

dt <- 0.01

TT <-250

lam <-2.75

sigma <-9.74*sqrt(dt)

n <-17.69

nn<-n

tt <- 0.0

for(i in 1:TT){

dn <-(lam*n)*dt+sigma*rnorm(1)

n <-dn+n

if(i %% 1==0){

nn<-rbind(nn,n)

tt<-rbind(tt,i);

}

}

matplot(tt,nn,type="l",ylab="Number of Cancer cells", xlab="Time",col="black",ylim=c(0,1e+05))

}

\###

\# jmmsd26ken()

\###

jmmcd26ken <-function(){

dt <- 0.01

TT <-250

lam <-2.75

sigma <-9.74*sqrt(dt)

n <-17.69

nn<-n

tt <- 0.0

for(i in 1:TT){

dn <-(lam*n)*dt-sigma*rnorm(1)

n <-dn+n

if(i %% 1==0){

nn<-rbind(nn,n)

tt<-rbind(tt,i);

}

}

matplot(tt,nn,type="l",ylab="Number of Cancer cells", xlab="Time",col="black",ylim=c(0,1e+05))

}

\###

\# jmmsd26plot()

\###

jmmcd26plot <-function(){

jmmcd26ken()

par(new=TRUE)

jmmcd26ryu()

}

\# **Codes of Monte Carlo simulations. These codes generate the plot x times that calculates the above programming.**

\###
\# jmmcvgraph (x)

#Generating the function that draws jmmcvplot () x times in one sheet.

\###
jmmcvgraph<-function(x){
if(x<=0){
print("Please enter one or more.")
}else{
for(i in 1:x){
if(i>1){
par(new=TRUE)
jmmcvplot ()
}else{
if(i==1){
jmmcvplot ()
}
}
}
}
}

\###
\# jmmcd24graph (x)

\###
jmmcd24graph<-function(x){
if(x<=0){
print("Please enter one or more.")
}else{
for(i in 1:x){
if(i>1){
par(new=TRUE)
jmmcd24plot ()
}else{
if(i==1){
jmmcd24plot ()
}
}
}
}
}

\###
\# jmmcd26graph (x)

\###
jmmcd26graph<-function(x){
if(x<=0){
print("Please enter one or more.")
}else{
for(i in 1:x){
if(i>1){
par(new=TRUE)
jmmcd26plot ()
}else{
if(i==1){
jmmcd26plot ()
}
}
}
}
}
==========

**Fig 2(a) and Fig 3(a) code**

#Note: This graph is written by Python, matplotlib

import numpy as np

import matplotlib.pyplot as plt

JMMcvm = (1., 8.93, 23.46, 79.63)

JMMcvsd = (0., 0.15, 2.79, 57.90)

JMMsh24m = (1.,7.37, 13.47, 29.18)

JMMsh24sd =(0., 0.28, 0.43, 18.91)

JMMsh26m = (1., 4.40, 10.44, 24.04)

JMMsh26sd =(0., 0.87, 2.29,4.57)

#the above six results are same as those listed in Table 1

n_groups = 4

fig, ax = plt.subplots()

index = np.arange(n_groups)

bar_width = 0.3

\# Drawing the bar graph

JMMcvBar = plt.bar(index, JMMcvm, bar_width,

alpha=opacity,

color='blue',

yerr=JMMcvsd,

error_kw=error_config)

JMMsh24Bar =plt.bar(index +bar_width, JMMsd24m, bar_width,

alpha=opacity,

color='red',

yerr=JMMsh24sd,

error_kw=error_config)

JMMsh26Bar = plt.bar(index +2*bar_width, JMMsh26m, bar_width,

alpha=opacity,

color='black',

yerr=JMMsh26sd,

error_kw=error_config)

plt.xlabel('Time')

plt.ylabel('Number of cancer cells')

plt.xticks(index + 2*bar_width / 3, ('Day1', 'Day4', 'Day7', 'Day10'))

plt.tight_layout()

plt.show()

**Fig 4 code**

\# Simulation code of $n_{0}=1 and \lambda=2.5$

\###

\# ryu pathway

\##

ryu <-function(){

dt <-0.01

TT <-250

lam <-2.5

sigma <-sqrt(lam*1*2)*sqrt(dt)

n <-1

nn<-n

tt <- 0.0

for(i in 1:TT){

dn <-(lam*n)*dt+sigma*rnorm(1)

n <-dn+n

if(i %% 1==0){

nn<-rbind(nn,n)

tt<-rbind(tt,i);

}

}

matplot(tt,nn,type="l",ylab="Number of Cancer cells", xlab="Time",col="black", ylim=c(0,750))

}

\##

#ken pathway

\##

ken <-function(){

dt <-0.01

TT <-250

lam <-2.5

sigma <-sqrt(lam*1*2)*sqrt(dt)

n <-1

nn<-n

tt <- 0.0

for(i in 1:TT){

dn <-(lam*n)*dt-sigma*rnorm(1)

n <-dn+n

if(i %% 1==0){

nn<-rbind(nn,n)

tt<-rbind(tt,i);

}

}

matplot(tt,nn,type="l",ylab="Number of Cancer cells", xlab="Time",col="red", ylim=c(0,750))

}

\##

\# Simulation code of growing cancer clones

\##

#Ryu code of the JMM CV cell lines

jmmngryucv <-function(){

dt <- 0.01

TT <-250

lam <-2.44

sigma <-18.73*sqrt(dt)

n <-188.87

nn<-n

tt <- 0.0

for(i in 1:TT){

dn <- sigma*rnorm(1)

n <-dn+n

if(i %% 1==0){

nn<-rbind(nn,n)

tt<-rbind(tt,i);

}

}

matplot(tt,nn,type="l",ylab="Number of Growing clones", xlab="Time", col="black", ylim=c(0,250))

}

#Ken code of the JMM CV cell lines

jmmngkencv <-function(){

dt <- 0.01

TT <-250

lam <-2.44

sigma <-18.73*sqrt(dt)

n <-188.87

nn<-n

tt <- 0.0

for(i in 1:TT){

dn <- -sigma*rnorm(1)

n <-dn+n

if(i %% 1==0){

nn<-rbind(nn,n)

tt<-rbind(tt,i);

}

}

matplot(tt,nn,type="l",ylab="Number of Growing clones", xlab="Time", col="red", ylim=c(0,250))

}

\# JMM sh-cd26 simulations

\# Ryu code

jmmcd26ngryu <-function(){

dt <- 0.01

TT <-250

lam <-2.75

sigma <-9.74*sqrt(dt)

n <-17.69

nn<-n

tt <- 0.0

for(i in 1:TT){

dn <-sigma*rnorm(1)

n <-dn+n

if(i %% 1==0){

nn<-rbind(nn,n)

tt<-rbind(tt,i);

}

}

matplot(tt,nn,type="l",ylab="Number of Growing Cancer cells", xlab="Time", col="black", ylim=c(-10,30))

par(new=TRUE)

matplot(tt, 0*tt, type="l", lty=6, ylab="Number of Growing Cancer cells", xlab="Time", col="blue", ylim=c(-10,30))

}

#Ken code

jmmcd26ngken <-function(){

dt <- 0.01

TT <-250

lam <-2.75

sigma <-9.74*sqrt(dt)

n <-17.69

nn<-n

tt <- 0.0

for(i in 1:TT){

dn <-sigma*rnorm(1)

n <-dn+n

if(i %% 1==0){

nn<-rbind(nn,n)

tt<-rbind(tt,i);

}

}

matplot(tt,nn,type="l",ylab="Number of Growing Cancer cells", xlab="Time", col="red", ylim=c(-10,30))

par(new=TRUE)

matplot(tt, 0*tt, type="l", lty=6, ylab="Number of Growing Cancer cells", xlab="Time", col="red", ylim=c(-10,30))

}

\# Plotting the growing clones of the JMM sh-CD26 cell line

jmmcd26ngplt <- function(){

jmmcd26ngken()

par(new=TRUE)

jmmcd26ngryu()

}
